# The RNA helicase eIF4A as a novel target in insect cells to combat arboviral infections

**DOI:** 10.1101/2025.10.22.683850

**Authors:** Tanja Rehling, Kim Mentchen, Leonie Konopka, Wiebke Obermann, Friedemann Weber, Patrick Schmerer, Marc F. Schetelig, Arnold Grünweller, Irina Häcker, Francesca Magari

**Affiliations:** Department of Insect Biotechnology in Plant Protection, Justus Liebig University Giessen, 35394 Giessen, Germany; Liebig Centre for Agroecology and Climate Impact Research, International Atomic Energy Agency Collaborating Centre, Justus Liebig University Giessen, 35394 Giessen, Germany; Institute of Pharmaceutical Chemistry, Philipps University Marburg, 35032 Marburg. Germany; Institute for Virology, FB10-Veterinary Medicine, Justus-Liebig University, 35392 Giessen, Germany

**Keywords:** *Aedes aegypti*, arboviruses, eIF4A, rocaglates, pateamines, antiviral, translation initiation, Rift Valley fever virus

## Abstract

Arboviruses transmitted by mosquitoes cause major global health burdens, yet few vaccines or antivirals exist. Targeting host factors required for viral replication offers a promising approach. The DEAD-box RNA helicase eIF4A, a core component of the translation initiation complex eIF4F, unwinds structured 5′ UTRs and is critical for many viral RNAs. We tested whether the natural products rocaglates and pateamines, potent eIF4A inhibitors in mammalian cells, also suppress arboviral replication and translation in insect vectors. Silvestrol strongly inhibited Rift Valley fever virus (RVFV) replication in human A549 cells without cytotoxicity, expanding the list of arboviruses dependent on eIF4A. Sequence analysis showed conservation of the rocaglate-binding motif between the arboviral vector *Aedes aegypti* and the fruit flies *Anastrepha suspensa* and *Drosophila melanogaster*. Dual-luciferase assays in insect cells of all three species confirmed silvestrol selectively inhibited translation from purine-rich reporters below cytotoxic thresholds. Purified eIF4A variants retained helicase activity, allowing direct testing of inhibitor interactions. Thermal shift assays demonstrated robust stabilization of eIF4A–RNA complexes by both compound classes in the wildtype proteins, with unexpected rocaglate sensitivity of the putatively insensitive *Ae. aegypti* H161L mutant, indicating a unique binding pocket geometry of the mosquito protein. Our results identify eIF4A as a conserved and druggable host factor in insects, highlighting its potential as a novel target for transmission-blocking antivirals and insect-specific inhibitors.

**Author Summary:** Natural compounds from the classes of rocaglates and pateamines have shown their potential to inhibit tumor cell growth and to effectively suppress replication of different viruses, including Ebola, corona, picorna, and arboviruses by inhibiting translation initiation through blocking of one of the initiation actors, the helicase eIF4A. eIF4A is needed to unwind structured 5’UTRs of selected mRNAs to allow binding of the 43S preinitiation complex. The compounds bind in a pocket of eIF4A, thereby clamping the RNA onto the eIF4A surface. Although eIF4A is highly conserved in eukaryotes not all eIF4A are sensitive to rocaglates. Mainly one key amino acid in the binding pocket determines sensitivity or insensitivity. In this study we expanded the list of arboviruses as potential targets of rocaglate-based drugs. Moreover, we approached the inhibition of arboviral infections from the vector side by studying the interaction between rocaglates and pateamines with eIF4A from the yellow fever mosquito *Aedes aegypti*, one of the most important arboviral vectors. Our study shows that also the mosquito eIF4A can be targeted by both compound classes. Moreover, our results reveal a unique architecture of the binding pocket that retains sensitivity even upon mutation of the key amino acid.

## Introduction

Arthropod-borne viruses (arboviruses) transmitted by mosquitoes, ticks, and sandflies cause substantial global morbidity. Major mosquito-borne examples, Zika, Dengue, Chikungunya, Rift Valley fever, West Nile, and Yellow fever, are frequently vectored by *Culex* and *Aedes* spp. (1). Disease reduction can target the vector or the virus. Current vector control relies heavily on insecticides, while vaccines and antivirals remain limited. Thus, sustainable vector control and broad-spectrum antivirals are urgently needed.

Targeting host factors essential for viral replication can provide pan-antiviral activity. A key host factor is the eukaryotic translation initiation factor 4A (eIF4A), a DEAD-box RNA helicase within the eIF4F complex that resolves structured 5′ UTRs to enable 43S preinitiation complex loading (2). Many viral RNAs contain structured 5′ UTRs and depend on eIF4A, making it an attractive pan-antiviral target (3–10). eIF4A inhibitors have shown activity against diverse human pathogens, including several arboviruses, in vitro, ex vivo, and in vivo (11–14). Such inhibitors could also be deployed in vectors or during transmission (e.g., topical formulations), potentially reducing host toxicity (15, 16).

Rocaglates and pateamines are two natural compound classes (Fig. S1) that inhibit eIF4A via RNA clamping: They stabilize eIF4A on purine-rich sequences and block unwinding, preventing 43S recruitment (17, 18). Structural work with human eIF4A, RocA, and a (AG)₅ RNA revealed π–π stacking with Phe163 and purines A7/ G8, and hydrogen bonds involving Gln195 and G8 (Fig. S2) (9, 19). Silvestrol’s dioxanyloxy moiety engages an arginine-rich pocket critical for complex formation (Fig. S1 and S2) (14) and can broaden sequence tolerance via interaction with a third nucleotide (A9) (20). Amino acid 163 (human numbering) is key for rocaglate sensitivity, F/Y/H support binding, whereas L/G/S confer insensitivity (14, 21). In contrast, pateamine-mediated clamping is largely independent of this position (20, 21).

So far, we and others have shown broad and potent antiviral activity of eIF4A inhibitors against a large set of highly pathogenic viruses such as Ebola-, Lassa-, Crimean Congo hemorrhagic fever-, and coronavirus, including the arboviruses Zika and Chikungunya *in vitro*, *ex vivo*, and also *in vivo* (*3–6, 8, 10*). Here we extend eIF4A inhibitor activity to Rift Valley fever virus (RVFV) and test whether rocaglates and pateamines inhibit translation also in insect vector cells. We compare eIF4A sensitivity across *Aedes aegypti*, *Anastrepha suspensa*, and *Drosophila melanogaster* using cell-based assays and purified wild-type and mutant proteins. We show robust eIF4A–RNA clamping by both inhibitor classes, species-specific differences in rocaglate sensitivity likely due to binding pocket variation and provide a basis for designing insect- targeted eIF4A inhibitors.

## Results

### Silvestrol inhibits RVFV replication

Rocaglate sensitivity has been shown for multiple arboviruses (9). Here, we tested Rift Valley fever virus (RVFV; family *Phenuiviridae*, order *Bunyavirales*) using two strains (MP-12 and Clone 13). A549 cells pretreated with 50 nM silvestrol and infected at MOI 0.1 showed a strong reduction of RVFV L-segment RNA for both strains relative to DMSO controls (Fig. **1a,b**). No cytotoxicity was detected at 50–100 nM (Fig. **1c**). These data indicate RVFV replication is sensitive to eIF4A inhibition and consistent with eIF4A-dependent viral protein synthesis.

**Fig 1:**
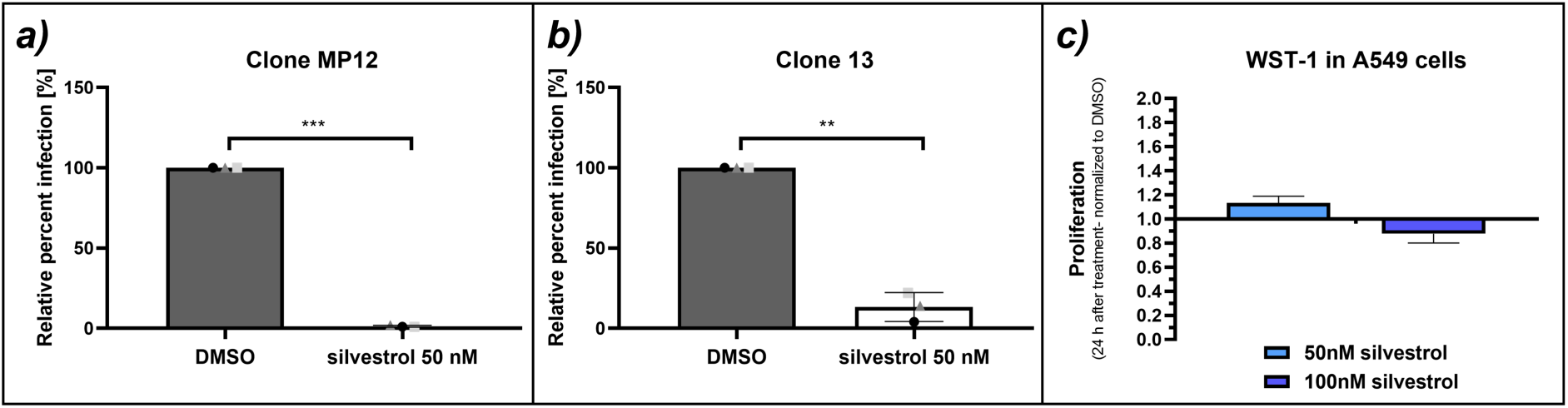
Effect of silvestrol on RVFV. RVFV L-segment RNA in A549 cells infected with (a) MP-12 or (b) Clone 13 after pretreatment with 50 nM silvestrol; (c) WST-1 assay in A549 cells (50–100 nM) to assess cytotoxicity. * p < 0.05; ** p < 0.01 and *** p ≤ 0.001.

### Conservation of amino acids in the eIF4A RNA binding pocket between human and insects

Instead of or in addition to targeting the arbovirus by rocaglates or pateamines, the vector itself could also be a target. To estimate if known human eIF4A inhibitors may also have an effect on one of the main arboviral vectors, the similarity between eIF4A from the yellow fever mosquito *Ae. aegypti* (Culicidae) and human eIF4A was analyzed. In addition, we were interested in the sensitivity of other insect families from the order of Diptera and chose *A. suspensa* as a representative for the Tephritidae and *D. melanogaster* of the Drosophilidae.

At the molecular level, inhibition of eIF4A by RNA-clamping is mediated by interactions of rocaglates with at least two purines in the bound RNA substrate and π-π stacking interactions with a phenylalanine (F163) as part of a six amino acid motif in the rocaglate binding pocket of the *human* eIF4A. Comparison of the amino acid sequence in the RNA binding pocket of eIF4A showed that the six amino acid motif is highly conserved between *Homo sapiens* and the insect species under investigation (Figure 2). Only at position 3 of the motif, corresponding to human Phe163, variation was observed. However, the amino acid substitutions found in the insect proteins (His or Tyr) all preserve RNA-clamping ability of rocaglates (14). We, therefore, predicted that a polypurine sequence should be clamped on the surface of eIF4A from these organisms. Instead, a leucine, isoleucine, serine, or glycine at this position should prevent rocaglate binding (14, 21). Interestingly, RNA-clamping of pateamines is independent from the amino acid at position 163 in human eIF4A (21).

**Fig 2:**
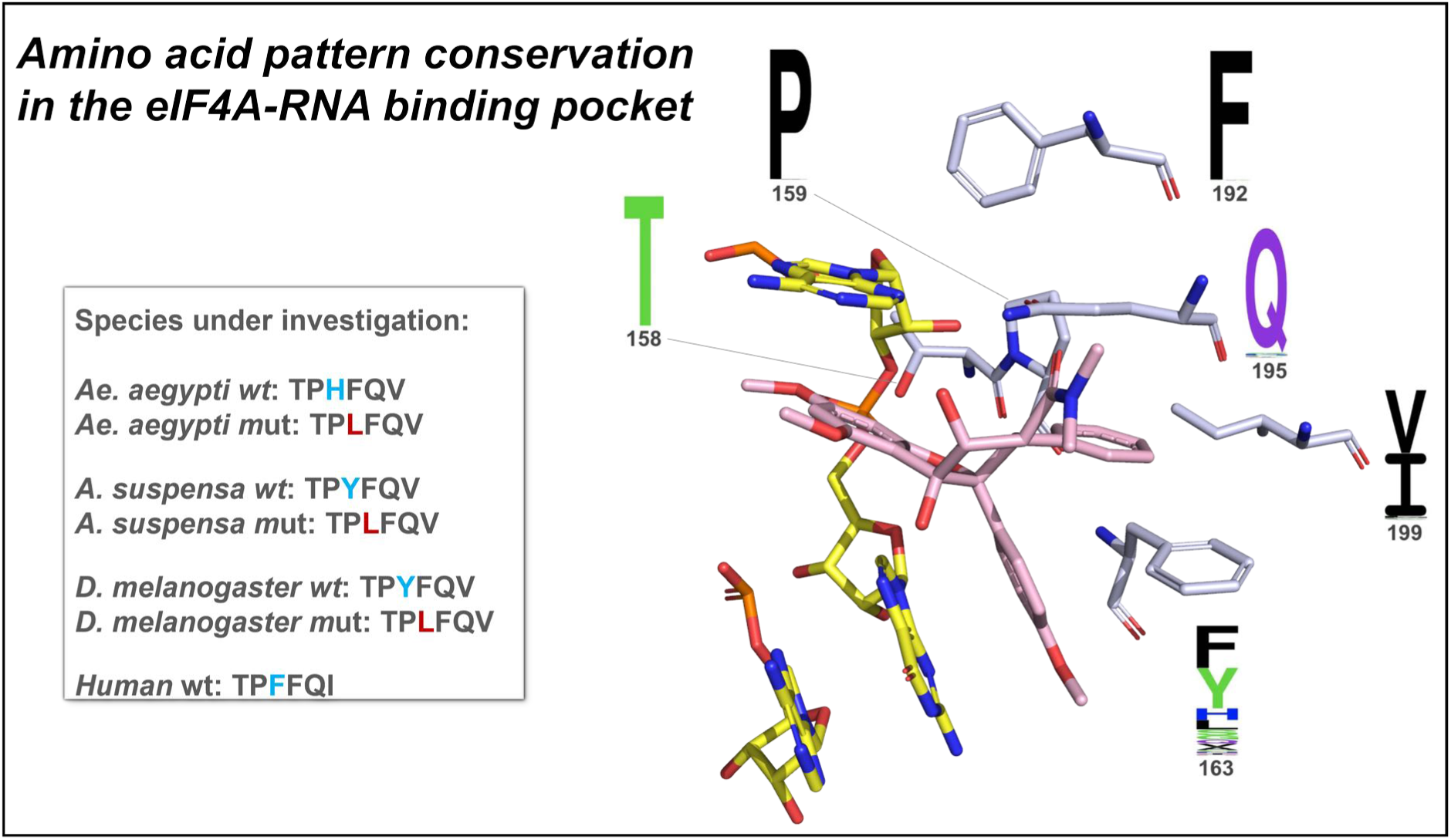
Comparison of amino acid pattern in the eIF4A-RNA binding pocket between *human* and flies species under investigation. Right: RocA (pink) bound to human eIF4A with (AG)₅ RNA (PDB: 5ZC9), highlighting key residues (T158, P159, F163, F192, Q195, I199). Left: six-residue motif across species; human F163 (light blue) corresponds to H161 (*Ae. aegypti*) and Y160 (*A. suspensa*, *D. melanogaster*). Mutations introducing L (red) model rocaglate insensitivity.

### Silvestrol inhibits translation initiation in insect cells

As the amino acid sequence in the insect eIF4A RNA binding pockets indicated rocaglate sensitivity, we tested the effects of silvestrol on translation efficiency in Aag2 (*Ae. aegypti*) and S2 (*D. melanogaster*) cell lines using a modified dual luciferase reporter assay (DLA). DLA plasmids contained a polypurine sequence (AG)_15_ upstream of the firefly luciferase CDS, which enables RNA clamping by rocaglates. In case of sensitivity, translation of firefly luciferase should therefore be inhibited in the presence of silvestrol. A control plasmid contained instead a mixed purine-pyrimidine sequence (AC)_15_, which confers eIF4A independence (7, 19). The original DLA plasmids, designed for use in human cells (6), were adapted for use in insect cells by exchanging the promoter and HCV IRES element (Figure S3).

To exclude the possibility that the reduction in luciferase expression is due to the use of cytotoxic concentrations of silvestrol in the DLA, instead of a specific interaction between silvestrol and the eIF4A-RNA complex, cytotoxicity was first re-evaluated for Aag2 and S2 cells using the WST-1 cell proliferation assay. Different sensitivities to silvestrol were observed: while the CC_50_ of the *Aedes* Aag2 cell line was 55 nM after 24 h incubation and 45.4 nM after 48 h, *Drosophila* S2 cells exhibited higher sensitivity, with a CC_50_ value of 11.8 nM after 24 hours of incubation and 7.1 nM after 48 hours (Fig. 3). Based on these results, silvestrol concentrations were titrated in the DLA up to the CC_50_ values.

**Fig 3:**
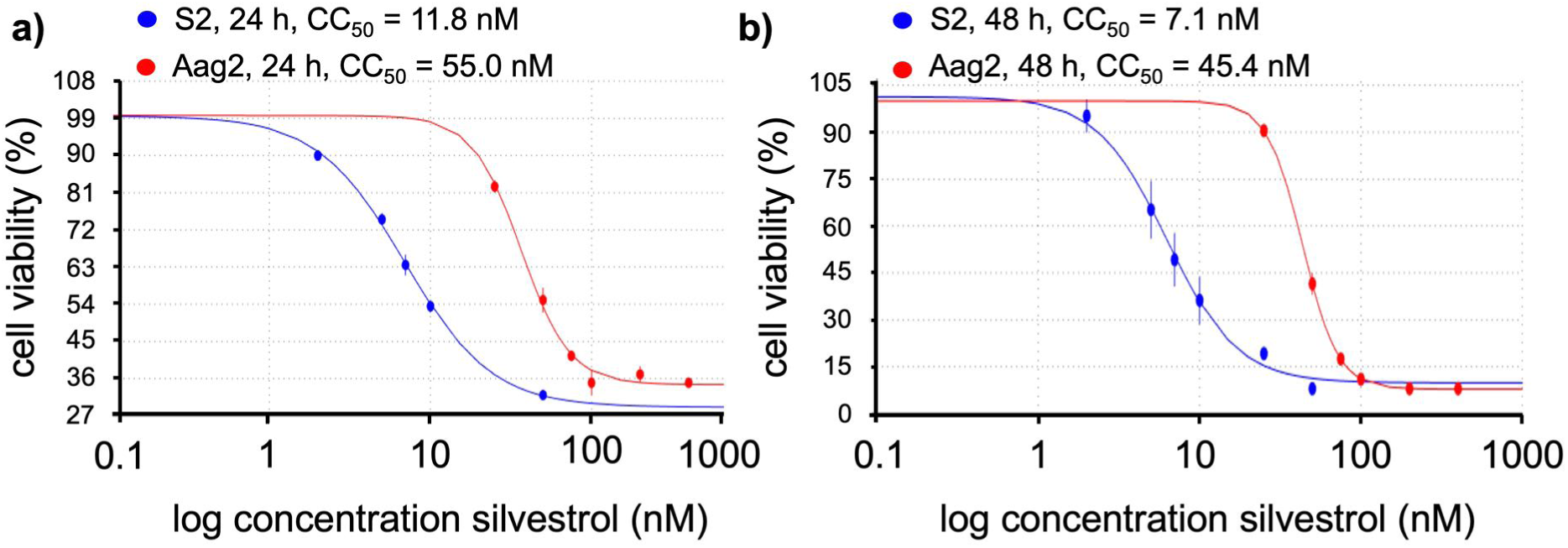
CC_50_ value determination for silvestrol cytotoxicity in Aag2 and S2 cells. a) incubation in the presence of silvestrol for 24 h, b) incubation for 48 h. CC_50_ was calculated using the LC_50_ Calculator software (22) and nonlinear regression analysis. Data shown are based on 2-5 replicates, except for Aag2 0 nM and 400 nM and S2 0 nM and 50 nM (24 h) and 50 nM (48 h), which were only determined once.

For both cell lines, silvestrol concentrations in the sub-CC_50_ range resulted in a strong reduction in firefly luciferase activity in the presence of the (AG)_15_ sequence upstream of the luciferase CDS, but not with the control plasmid (AC)_15_ (Fig. 4), indicating efficient purine-dependent RNA-clamping by silvestrol onto eIF4A. Silvestrol concentrations in the CC_50_ range led to a reduction of translation efficiency, also with the (AC)_15_ control, confirming the cytotoxicity results of the cell proliferation assays. In Aag2 cells, this effect was already visible at the half CC_50_ concentration (Fig. 4b), potentially reflecting added stress to the cells due to the transfection, or an increased silvestrol uptake mediated by the transfection reagent.

**Fig 4:**
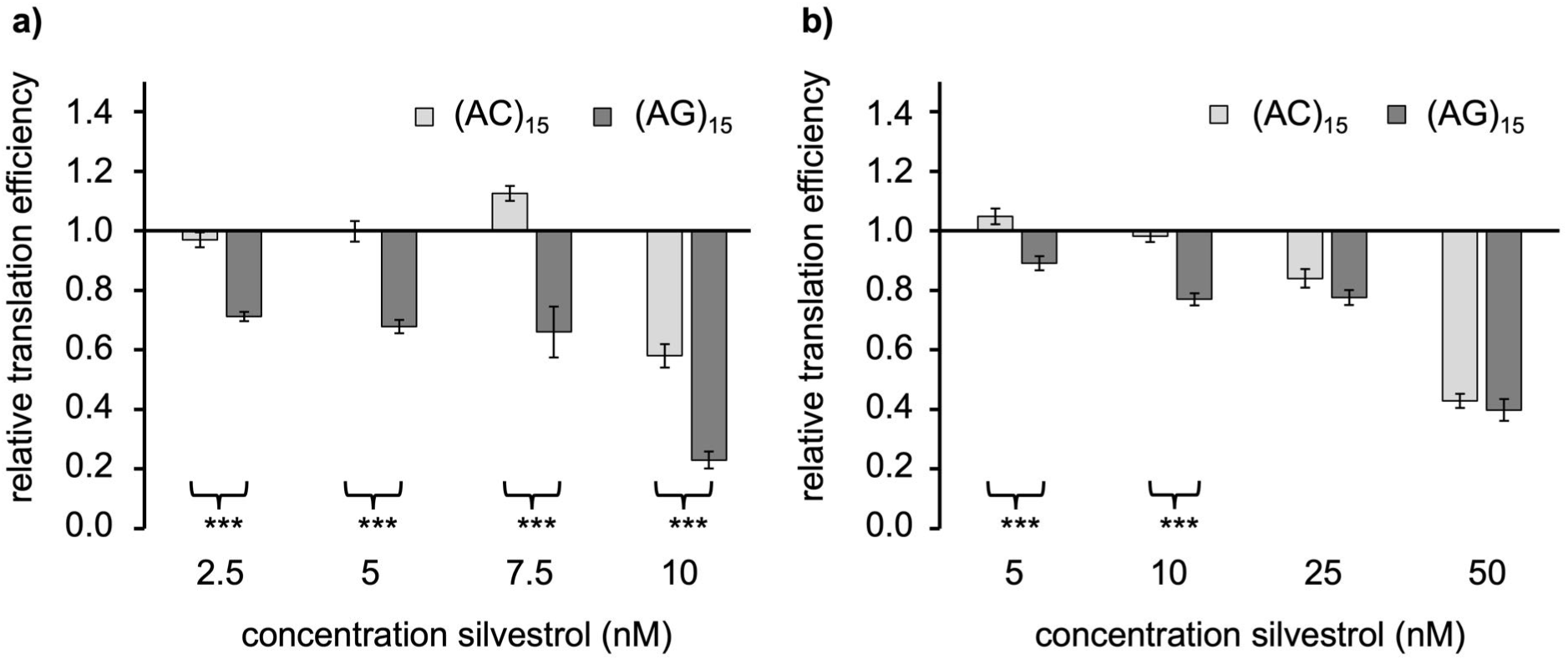
Effect of silvestrol on translation efficiency determined by DLA in a) *D. melanogaster* S2 cells (t = 24 h) and b) *Ae. aegypti* Aag2 cells (t = 48 h). Interaction of silvestrol with insect eIF4A was tested with a polypurine (AG)₁₅ in the 5′ UTR of the reporter construct. The same construct with a mixed purine-pyrimidine (AC)₁₅ sequence served as silvestrol- insensitive control. Shown are means of the relative translation efficiency (firefly activity normalized to renilla activity and to the corresponding DMSO controls). Error bars represent SEM. Data shown are based on 2-3 biological replicates with 3 technical replicates each, with the exception of the 50 nM experiment, which was performed only once; * p < 0.05; ** p < 0.01; *** p ≤ 0.001.

### Overexpressed insect eIF4A variants show unwinding activity

After demonstrating that our inhibitors reduce translation in insect cells, we speculated that mutants with a Leu instead of a His or Tyr (Fig. 2) should not be capable of rocaglate-mediated RNA clamping, while pateamines should have the potential to clamp RNA independently of the amino acid at this position. To prove our assumption, we first overexpressed and purified the insect eIF4A wt and mutant (with Leu) variants (Fig. 5 a-c) and tested the functionality of the purified proteins in a previously described helicase assay (21). As shown in Fig. 5 d-f, all purified eIF4A enzymes showed unwinding activity, demonstrating that the expressed wt as well as mutant versions of eIF4A were functional.

**Fig 5:**
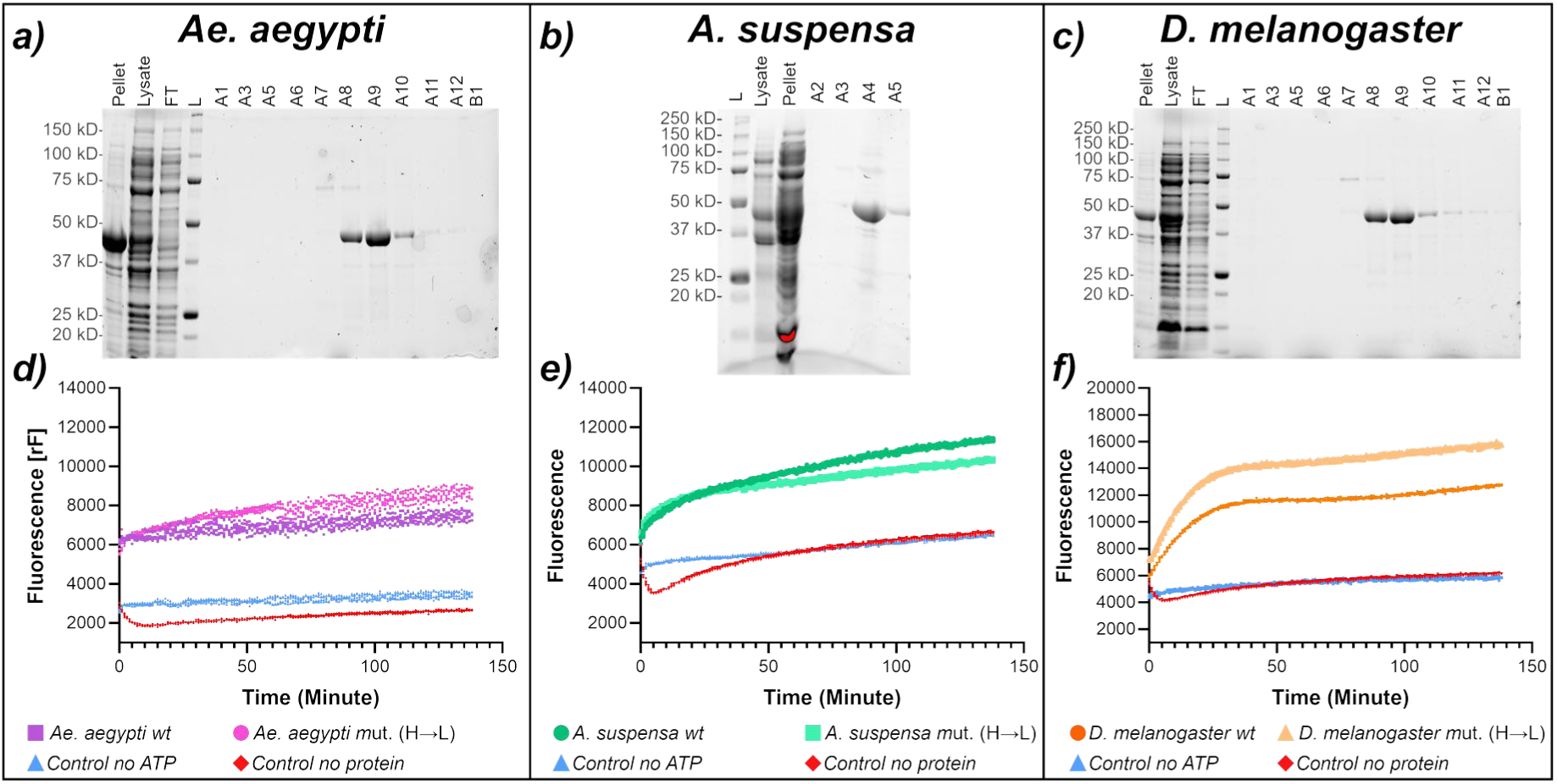
Protein purification and helicase assays. a) – c) TGX stain-free PAGE images showing successful protein expression and purification of eIF4A wt from a) *Ae. aegypti* (MW = 47.6 kDa), b) *A. suspensa* (MW = 46.0 kDa) and c) *D. melanogaster* (MW = 48.0 kDa). From left to right, the cell pellet, lysate, flow-through (FT), the leader Precision Plus Protein^TM^ Unstained Protein Standards (10 - 250 kDa) (L), and the elution fractions; collected were fractions A8-10 in a), A4-5 in b), and A8-10 in c). d) – f) helicase assays showing the increase in fluorescence over time through eIF4A-mediated unwinding of quenched, fluorescently labelled RNA substrates; d) *Ae. aegypti* (wt curve in purple, mutant H161L in pink), e) *A. suspensa* (wt curve in dark green, mutant Y160L in light green), and c) *D. melanogaster* (wt curve in dark orange, mutant Y160L in pale orange). Reactions without ATP (light blue) and without protein (red) served as negative controls. Data shown are based on at least three replicate experiments.

### Interaction of purified insect eIF4A with rocaglates and pateamines

Next, we tested in thermal shift assays (TSA) if the purified eIF4A enzymes in complex with the polypurine sequence (AG)_5_ can interact with different rocaglates (CR-31-B (-), RocA, silvestrol, zotatifin) and pateamines (PatA, desmethyl-desamino PatA). As a negative control, we used the non-functional enantiomer of CR-31-B (-), namely, CR-31-B (+) (7). As expected, both compound classes strongly increased the thermal stability of wt eIF4A-(AG)_5_ complexes by mostly 5 to 10 °C (Fig. 6a-c). The overall strongest effect was observed for PatA with *Ae. aegypti* wt protein with more than 13 °C increase, whereas the weakest effect was observed for zotatifin with *D. melanogaster* eIF4A showing only an increase of 3 °C in thermal stability (Fig. 6a, c).

**Fig 6:**
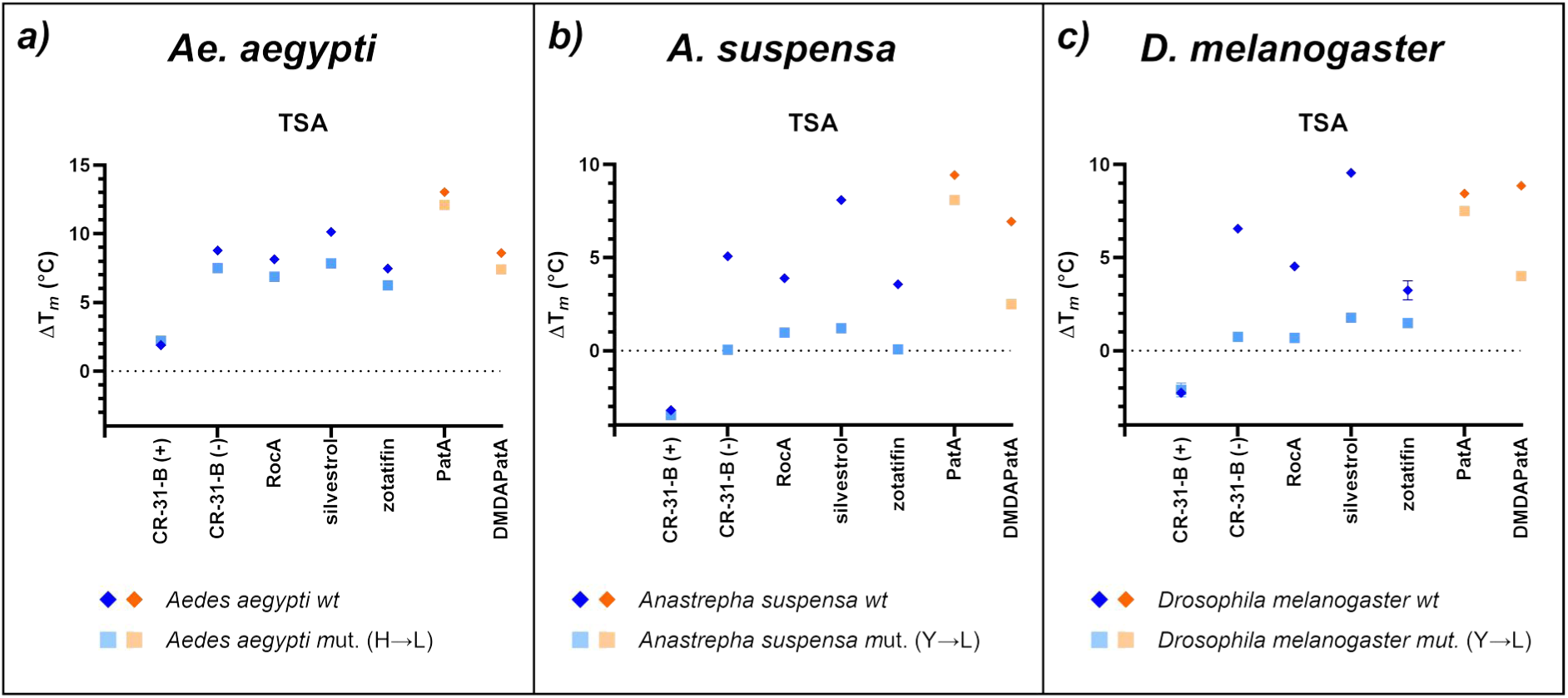
Effect of different rocaglates and pateamines on eIF4A-polypurine complex stability with wt and mutant protein versions. TSA of eIF4A from a) *Ae. aegypti*, b) *A. suspensa*, and c) *D. melanogaster* to evaluate the clamping of rocaglates (blue) and pateamines (orange) to the eIF4A-RNA complex. The wt proteins are shown in dark blue/orange rhombuses and the mutants (mut.) in light blue/orange squares. The difference in melting temperature ΔT*_m_* (°C) was calculated between eIF4A and the eIF4A-(AG)_5_ -AMP-PNP complex with different inhibitors. Data shown are based on at least three replicate experiments. ΔT*_m_* (°C) with the corresponding standard error of the mean (SEM) for n ≥ 3 is shown in the supplementary Table S1.

Interestingly, silvestrol with its additional dioxan moiety has the strongest thermal stabilization effect in all three eIF4A variants from flies compared to the other rocaglates that lack the dioxan ring. As predicted, the mutant versions of eIF4A from *A. suspensa* and *D. melanogaster* showed no significant increase in thermal stability with rocaglates, indicating that these molecules cannot clamp the RNA when a Leu is present instead of Tyr. Unexpectedly, the situation in *Ae. aegypti* was different since the H to L mutant showed similar increase in thermal stability as the wt (Fig. 6a, light and dark blue), indicating that in this mosquito the pocket for binding of rocaglates must be arranged in an unexpectedly different manner (s. Discussion).

For the pateamines, PatA was able to clamp (AG)_5_ also in the mutants as expected (Fig. 6a-c, red and orange), and showed a stronger clamping effect with *Aedes* and *Anastrepha* proteins than the rocaglates. DMDAPatA, however, behaved differently. It showed a weaker thermal stabilization effect than PatA in *Ae. aegypti* and *A. suspensa* (Fig. 6a, b). Moreover, there was a strong difference in thermal stability between wt and mutant protein in *A. suspensa* and *D. melanogaster* (Fig. 6b, c).

### Different binding affinities for polypurines, purine-pyrimidine mixes, and polypyrimidines

In a next set of TSA experiments we tested which RNA substrates can form a complex with the eIF4A wt and mutant proteins (Fig. 7). Thermal stabilization was analyzed in the presence of silvestrol or CR-31-B (-) with different RNA-10mers as substrates. Silvestrol and CR-31-B (-) were selected due to their structure difference. The additional dioxane moiety of silvestrol may broaden the interactions with different RNA oligos. Substrates consisted of polypurines ((AG)_5_, (GA)_5_), mixed purine-pyrimidine sequences ((UG)_5_, (UA)_5_, (AC)_5_) or the polypyrimidine sequence (UC)_5_. In the presence of polypurines, we observed the expected increase in TSA for both compounds when eIF4A wt was used. The sequence (UG)_5_ produced a reduced thermal shift for *A. suspensa* and *Ae. aegypti*. However, in *D. melanogaster* the shift was comparable to that of polypurines. Changing the sequences to (UA)_5_, (AC)_5_ or to polypyrimidines prevented any thermal stabilization for all species (Fig. 7a, b). The Leu mutants of *A. suspensa* and *D. melanogaster* resulted in a strong decrease in thermal stabilization (dashed arrows in Fig. 7a, b), confirming the results from above. The *Ae. aegypti* mutant behaved again differently: in combination with (AG)_5_ there was no reduction of the thermal shift as observed before, whereas the (GA)_5_ 10mer showed a moderate but measurable thermal shift of about 5 °C with silvestrol and 3 °C with CR-31-B (-) compared to the wt protein (Fig. 7a, b). Again, this indicates that the mode of action of rocaglates in *Ae. aegypti* differs from other organisms in an unexpected manner. Overall, no substantial differences between silvestrol and the synthetic rocaglate CR-31-B (-) could be observed.

**Fig 7:**
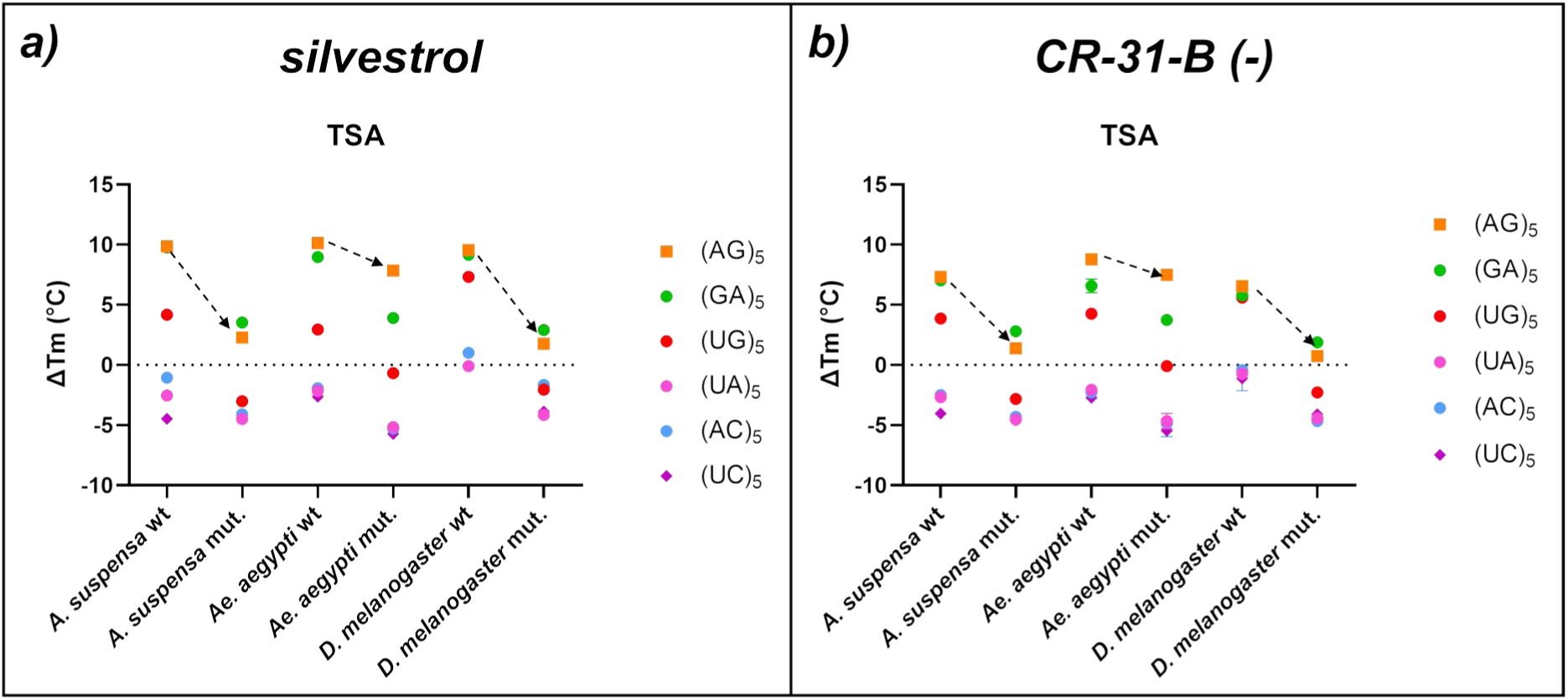
Effect of different RNA 10mers on rocaglate binding to eIF4A wt and mutant proteins. The rocaglates silvestrol (a) and CR-31-B (-) (b) were tested on polypurine stretches ((AG)_5_, orange and (GA)_5,_ green), a polypyrimidine stretch ((UC)_5_, violette) and mixed purine-pyrimidine stretches ((UG)_5_, red, (UA)_5_, pink, and (AC)_5_, light blue) in combination with *A. suspensa*, *Ae. aegypti* and *D. melanogaster* wt as well as the mutant (mut.) eIF4A versions. Data shown are based on at least three replicate experiments. The difference in melting temperature ΔT*_m_* (°C) was calculated between the eIF4A and the eIF4A-RNA-AMP-PNP complex with the two inhibitors. ΔT*_m_* (°C) with the corresponding standard error of the mean (SEM) for n ≥ 3 is shown in the supplementary Table S2.

## Discussion

The main goal of this study was to investigate whether known human eIF4A inhibitors can be used to target translation efficiency in insect cells as a strategy to block transmission of arboviruses and to compare the inhibitor sensitivity of an arboviral vector with insects from other families. In this study, we found for the first time that RVFV requires eIF4A for protein synthesis, and that eIF4As from three different insect families are sensitive to rocaglate and pateamine treatment.

By clamping eIF4A to selected mRNAs the rocaglates and pateamines negatively affect translation efficiency. A standard assay for measuring effects on translation efficiency in human cancer cell lines is the dual-luciferase reporter assay. By introducing a short polypurine sequence (AG)_15_ upstream of the firefly luciferase start codon, the effect of inhibitors on eIF4A- dependent translation can be investigated. Interestingly, the initial use of DLA plasmids designed for human cancer cells showed that neither the HSK promoter nor the HCV IRES element is functional in the insect cell lines used in this study. The reduction of the translation efficiency with the purine-containing 5’UTR but not with the purine-pyrimidine control in the presence of silvestrol indicated RNA-clamping by rocaglates onto the insect eIF4A’s with 5’UTRs that harbour polypurine stretches. The mode of action of eIF4A from flies compared to human eIF4A seems to be similar since also pateamines reduce translation efficiency. The observation that the faster proliferating S2 cells were more sensitive to silvestrol than the Aag2 cells also matched observations from human cell culture and primary cancer cells (23). While the S2 cells were more sensitive to silvestrol, no cytotoxicity was observed below the CC_50_, in contrast to Aag2 cells, which showed signs of unspecific cytotoxicity already at the half CC_50_ for silvestrol, possibly due to added stress to the cells by the transfection reagent or an increased cellular silvestrol uptake mediated by the reagent. Notably, the CC_50_ values for silvestrol in Aag2 and S2 cells determined here deviate approximately by a factor of 3 from those in previous studies with these cell lines (14).

Thermal shift assays with purified eIF4A wt proteins and mutants were used to prove the direct interaction of rocaglates or pateamines with eIF4A. While eIF4A wt and mutant proteins from *A. suspensa* and *D. melanogaster* behaved similarly to the *human* eIF4A with respect to rocaglate-sensitivity or insensitivity, eIF4A from *Ae. aegypti* showed an unexpected behavior. The H161L mutant was expected to be rocaglates-insensitive, because the Leu at this position (corresponding to *human* F163) should prevent clamping due to the loss of π-π stacking interactions between the aromatic ring of H161 and rocaglate rings B and C. Surprisingly, the *Ae. aegypti* H161L mutant still has the ability for RNA-clamping by rocaglates (Fig. 6a). This may be due to the different chemical and spatial properties of the amino acid residues surrounding the RNA binding pocket (Fig. 8). On the protein surface adjacent to the key amino acid F163 (H161 or Y160 in the studied insects) an asparagine residue is present in the human eIF4A (human N167) as well as in the one from *A suspensa* and *D. melanogaster* (N164), whereas *Ae. aegypti* has a serine residue (S165) at this position (Fig. 8b). Human N167 seems to close the rocaglate binding pocket, resulting in a narrower and deeper pocket with a more defined shape compared to that of *Ae. aegypti*. When F163 is mutated to L in the human eIF4A, the rocaglate ring C would clash with the latter (Fig. 8d), preventing the binding and the accommodation of the inhibitor in the binding pocket, probably also due to the closure of the pocket by N167. Accordingly, an L161 in *Ae. aegypti* clashes in the model with ring C of rocaglates. Binding may not be prevented in this case because the inhibitor has more space to accommodate, possibly due to the broader pocket created by S165. This may result in a conformational rearrangement of rocaglates in the RNA binding pocket of *Ae. aegypti,* that may not be possible in the *human* eIF4A due to N167 or N164 in *A. suspensa* and *D. melanogaster*. To clarify this, the exact mode of action of rocaglate binding in *Ae. Aegypti* should be addressed in a follow-up study.

**Fig 8:**
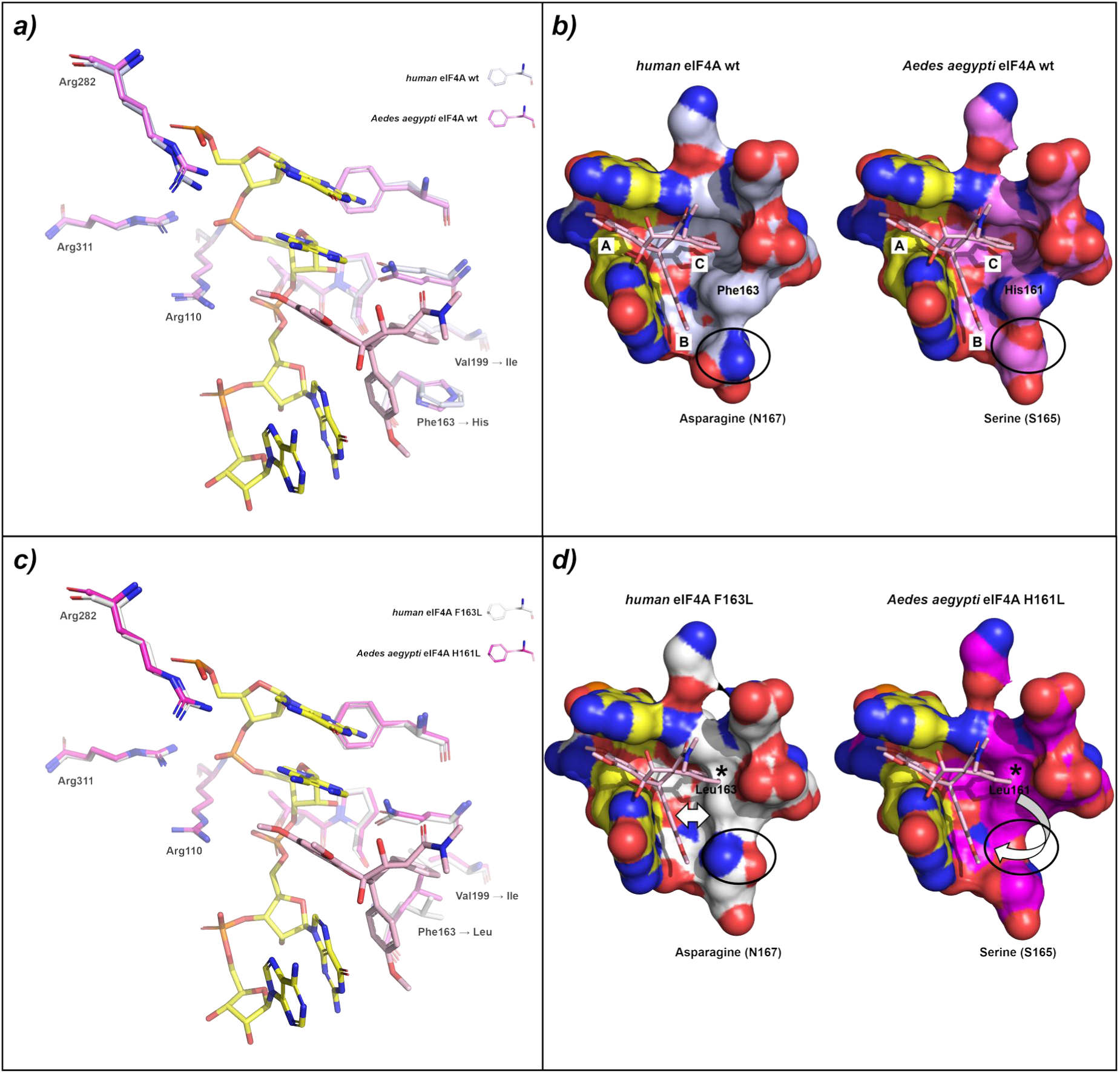
Alphafold3 structure comparison between a-b) *human* (silver) and *Ae. aegypti* (violet) eIF4A wt and c-d) *human* eIF4A H163L (white) and *Ae. aegypti* H161L (magenta) in complex with the polypurine (AG)_5_ and the inhibitor RocA. a) Typical amino acids pattern conservation in the eIF4A-RNA binding pocket as shown in Fig. 2 represented as stick model. The binding pockets of the *human* and *Ae. aegypti* eIF4A wt perfectly overlap. H161 in *Ae. aegypti* corresponding to F163 in *human* eIF4A preserves the typical π-π stacking interactions with RocA rings B and C (see Fig. S2a for more details), allowing the rocaglates to perfectly accommodate in the RNA binding pocket. b) Surface representation of a) highlighting the sub-pockets A, B and C as well as the key residues H161 and S165 in *Ae. aegypti* corresponding to F163 and N167 in *human* eIF4A. c) F163 in *human* eIF4A corresponding to H161 in *Ae. aegypti* has been mutated with a leucine residue and it is represented here as a stick model. Beside a different orientation of the leucine’s side chains, the other amino acids perfectly overlap as in case of a) d) Surface representation of the RNA binding pocket highlighting the clashes between ring C of RocA and the leucine residues (L163 in *human* and L161 in *Ae. aegypti*). In the *human* eIF4A an asparagine residue (N167) closes the RNA pocket whereas in the *Ae. aegypti the* corresponding serine residue (S165) renders the pocket broader. This may promote a conformational rearrangement and better accommodation of RocA in the RNA binding pocket of *Ae. aegypti* compared to the one of the *human* eIF4A. The combination of the clash with L163 and a smaller pocket may prevent RocA binding to the *human* eIF4A F163L-RNA complex. Contrarily, the clash with L161 of *Ae. aegypti* does not hamper the binding of RocA due to a higher freedom of movement in the broader RNA binding pocket.

## Materials and Methods

### Effect of silvestrol on RVFV infections

Human A549 cells were obtained from ATCC (Rockville, MD) and were pretreated with DMSO or 50 nM silvestrol for 1 h, triple-washed, and infected with (a) RVFV MP-12 or (b) RVFV Clone 13 at MOI 0.1. Cells were incubated in medium containing the corresponding compound concentration. Total RNA was extracted 18 h post-infection. RVFV L-segment RNA was quantified by two-step qRT-PCR and normalized to GAPDH as described (24). DMSO controls were set to 100%. Statistics: three independent experiments, paired two-sided t-test.

### Identification of insect eIF4A1 genes and cloning of variants for protein expression

Human eIF4A1 homologs were identified by BLAST in *D. melanogaster* (GCF_000001215.4, 2014) and *Ae. aegypti* (LVP AGWG L5.1, 2017). For *A. suspensa*, the homolog was inferred from *A. ludens* (GCF_028408465.1). The residue corresponding to human Phe163 is Tyr160 in *D. melanogaster* and *A. ludens* and His161 in *Ae. aegypti*.

Wild-type coding sequences were amplified from cDNA and cloned into pET- 28a(+)_eIF4A1(19–406) via Gibson Assembly using NdeI/XhoI-opened backbone (14): Primers P2192/P2132 (*D. melanogaster*) and P2127/P2128 (*Ae. aegypti*). *A. suspensa* eIF4A was first amplified (P2341/P2342), TOPO-cloned and sequenced (near-identity to *A. ludens*), then re-amplified (P2386/P2132) and inserted into pET-28a(+).

Site-directed mutagenesis introduced leucine at the key position (human F163 equivalents): *D. melanogaster* TAC→CTT (primers P2192/P2194 and P2193/P2132), *Ae. Aegypti* CAT→CTT (P2127/P2130 and P2129/P2128), *A. suspensa* TAC→CTT (P2386/P2385 and P2384/P2132). Overlap PCR products were Gibson-assembled into pET-28a(+). All PCRs used Q5 High-Fidelity DNA Polymerase (NEB). Primer sequences are in Table S3.

### Adapting Dual-luciferase vectors for testing the effects of silvestrol on translation in insect cells

The HSV-TK promoter pFR_HCV_xb_polyAC and pFR_HCV_xb_polyAG (8) was replaced with the hr5-ie1 enhancer/promoter, and the HCV IRES with the Drosophila C virus IRES. hr5- ie1 (25) was PCR-amplified from AH465 (pXLBacII_IE1hr5-DsRed.T3-SV40) (26) using P2108/P2109 (BglII/PstI overhangs) and ligated into BglII/PstI-digested vectors to yield V417 (polyAC) and V418 (polyAG) (Fig. S3). The DCV IRES was amplified from plasmid #707 (Rob Harrell, ITF) with P2317/P2318 and Gibson-cloned into SnaBI/NdeI-digested V417/V418 to give V428/V429. A shorter DCV IRES (27) was PCR-amplified from V429 (P2318/P2350, SpeI/NdeI) and ligated into SpeI/NdeI-digested V428/V429 to yield the final plasmids V432/V433.

### eIF4A protein variants expression and purification

The plasmids pET-28a(+)-His_6_-eIF4A1 from *Ae. aegypti*, *A. suspensa* and *D. melanogaster* (wt and mut.) were transformed into BL21 (DE3) competent cells. Pre-cultures were grown in 100 mL of LB media with kanamycin (50 μg/ml) at 37 °C, 150 rpm for 18h, after which 5 mL were transferred to 500 mL of LB media with kanamycin (50 μg/ml) and grown until the OD_600_ reached 0.5-0.6. Expression was induced with 0.5 mM Isopropyl β-D-1-thiogalactopyranoside (IPTG) overnight at 18 °C. Cells were pelleted at 10 000 rpm for 10 min at 4 °C, resuspended in Buffer A (20 mM HEPES–NaOH [pH 7.5], 300 mM KCl, 20 mM Imidazole, 0.1 mM EDTA, 10 mM β-mercaptoethanol and 10 % glycerol) and lysed by sonication (SONIFIER 250, BRANSON) in Buffer A containing lysozyme (50 µg/mL), benzonase (Benzonase® Nuclease, 5KU, Sigma-Aldrich/Merck) and one tablet of cOmplete™ Mini Protease Inhibitor Cocktail, EDTA-free (Roche. Germany). The lysates were centrifugated at 20 000 rpm for 45 min at 4 °C (Avanti™ J25. BECKMAN COULTER™) and the supernatant was collected and filtrated (Ultrafree®-MC filters, pore size 0.45 μm, Merck). The supernatants were loaded in 1 mL Ni-NTA column (HisTrap HP, GE Healthcare Life Science. Freiburg, Germany) and the column washed with 20 mL of Buffer A. Proteins eluted with 100 % of imidazole gradient at a flow rate of 1 mL/min in Buffer B (20 mM HEPES–NaOH [pH 7.5], 300 mM KCl, 800 mM Imidazole, 0.1 mM EDTA, 10 mM β-mercaptoethanol and 10 % glycerol). The eluted proteins were dialyzed overnight in the storage buffer (20 mM HEPES–NaOH [pH 7.5], 100 mM KCl, 5 mM MgCl_2_, 1 mM DTT and 10 % glycerol) and concentrated to 2 mg/mL. The purified protein was stored at −80 °C.

### Thermal Shift Assays

TSA experiments were performed on a real-time PCR system (QuantStudio™ 3, Applied Biosystems, Waltham, MA, USA) in a MicroAmp™ Fast Optical 96-well plate (Applied Biosystems, Waltham, MA, USA) using QuantStudio™ Design & Analysis software (version 1.4.2.). 5 μM of recombinant eIF4A was incubated with 50 μM of a polypurine RNA (AG)_5_ (Biomers, Ulm, Germany), 1 mM AMP-PNP (Roche, Basel, Switzerland), 100 μM of inhibitors (silvestrol, RocA, CR-31-B (+), CR-31-B (-), zotatifin, PatA, DMDAPatA) and 75 μM of SYPRO Orange (S6650, Invitrogen, Carlsbad, CA, USA) in 20 mM HEPES–KOH buffer pH 7.5, 100 mM KCl, 5 mM MgCl_2_, 1 mM DTT and 10 % (v/v) glycerol at RT. In the first step, the protein sample was subjected to a heating rate of 1.6 °C/s until a temperature of 10 °C was reached and kept constant for two minutes. In the second step, the temperature was increased by 0.05 °C/s until a temperature of 95 °C was reached and kept constant for one minute. In the final step, the temperature was decreased by 1.6 °C/s until 10 °C and kept constant for one minute. The wavelength of the fluorescence scan for excitation and emission was set to the spectroscopic maxima of SYPRO^®^ Orange (472 nm and 570 nm, respectively). The melting curves were analyzed using Protein Thermal Shift Software (version 1.3) from Thermo Fisher Scientific.

### eIF4A Helicase Assay

Two labeled RNAs were used: 10-mer-Cy3 (5′-[CY3]GCUUCCGGU-3′) and 16-mer-BHQ2 (5′- ACUAGCACCGGAAAGC[BHQ2]-3′), plus unlabeled 10-mer competitor (5′-GCU UUC CGGU- 3′). Duplexes were annealed (1 µM each; 80 °C 5 min, RT 1 h, ice 10 min). Reaction buffer: 150 mM HEPES–KOH pH 7.4, 15 mM Mg(OAc)₂, 10 mM DTT, 500 mM KOAc, 1 mM ATP.

Final: 100 nM RNA (ss or ds), 12.5 µM eIF4A. Controls lacked ATP or protein. Fluorescence was recorded on a Tecan Infinite M Plex.

### Insect cell culture

Aag2 cells (*Ae. aegypti*) (28), and S2 cells (*D. melanogaster)* (29) were cultured in complete Schneideŕs medium (Schneideŕs Drosophila Medium (Gibco) supplemented with 10 % fetal bovine serum (FBS, Merck-Sigma), 100 U/ml penicillin and 100 μg/ml streptomycin (Gibco) and 1 x MEM NEAA, Minimum Essential Medium (Gibco)) at 27 °C without CO_2_.

### Silvestrol toxicity assays

To determine the half-maximal cytotoxic concentration CC_50_ of silvestrol in the insect cell lines a proliferation assay was carried out. Cells were seeded at a density of 3 × 10⁴ cells per well (Aag2) or 2 × 10⁴ cells per well (S2) in a transparent 96-well culture plate with complete Schneideŕs medium and incubated for 24 hours at 27 °C to reach near confluency. Silvestrol or DMSO as control were added to a final concentration of 2.5 nM – 50 nM for S2 cells and 25 nM – 400 nM for Aag2 cells and cells were incubated at 27 °C. After 24 and 48 hours, cell proliferation was analysed using the WST-1 assay (Sigma Aldrich) following the manufacturer’s protocol. Absorbance was measured using a Tecan SPARK reader at 440 nm, with 650 nm as the reference wavelength, after two and three hours of incubation with the WST-1 reagent. All conditions were tested in triplicates. Percentage viability of the experimental groups in comparison to the control with 0 nM silvestrol / DMSO were computed and the CC_50_ value was calculated using non-linear regression with the “LC_50_ Calculator” software (22).

### Dual-luciferase reporter assays

Cells were seeded 24 h before transfection (Aag2: 3 × 10⁴; S2: 2 × 10⁴ cells/well) in black clear-bottom 96-well plates. Transfection used 90 ng plasmid/well and Lipofectamine 3000 in serum-free medium. Four hours later, medium was replaced with complete medium ± silvestrol / DMSO (S2: 2.5–10 nM, 24 h; Aag2: 5–50 nM, 48 h). Firefly and Renilla luciferase activities were measured with the Dual-Luciferase® Reporter Assay (Promega) using 50 µL LARII and Stop & Glo® per well on a Tecan SPARK. For S2, a filter reduced signal by 100× to avoid saturation. Firefly was normalized to Renilla and DMSO controls; SEMs were calculated on normalized ratios. Statistics: one-way ANOVA with Shapiro–Wilk and Brown–Forsythe tests; Holm–Šidák post hoc (30, 31). If variances were highly unequal or for unequal group sizes Kruskal–Wallis with Dunn’s test (32).

### Software

For protein purification, UNICORN^TM^ (Version 6.0) was used on an ÄKTA pure system (GE Healthcare). AlphaFold Server prediction was used to predict the structure of eIF4A variants from flies (Abramson et al. 2024). The PyMOL Molecular Graphics System (Version 2.0, Schrödinger, LLC) was used to both, analyze the predicted Alphafold3 structures and for picture generation. GraphPad Prism version 9 for Windows, (GraphPad Software, Boston, Massachusetts USA) was used for data and statistical analysis as well as to design the graph reported in the present work. CC_50_ values of silvestrol toxicity assays were calculated ’with the “LC_50_ Calculator” software (22), Statistcs analysis of the dual-luciferase reporter assays was performed with SigmaPlot (Systat Software GmbH).

## Acknowledgements

We thank Robert Harrell, Insect Transformation Facility, University of Maryland, MD, USA, for providing plasmid #707 (Drosophila C virus IRES).

We thank Prof. Dr. Alois Fürstner from the Max-Planck-Institut für Kohlenforschung (Mülheim Ruhr, Germany) for providing us with pateamines.

## Supplementary Information

**Table S1: TSA values of eIF4A variants against rocaglates and pateamines in complex with (AG)_5_**

**Table S2: TSA values of eIF4A variants in complex with different RNA oligos**

**Table S3: Primers**

**Table S4: Sequences**

**Figure S1: Chemical structure of eIF4A inhibitors.** On the top: the natural rocaglates rocaglamide A (RocA) and silvestrol isolated from the plant *Aglaia foveolate* and the natural pateamine pateamine A (PatA) isolated from the marine sponge *Mycale hentscheli*. The characteristic cyclopenta[b]benzofuran ring system of rocaglates is highlighted in green and rings A, B and C which address the respective pockets A, B and C in the eIF4A binding pocket are labelled in the RocA structure. On the bottom: the synthetic rocaglates CR-31-B (-), CR- 31-B (+) and zotatifin and the synthetic pateamine desmethyl desamino pateamine A (DMDA- PatA).

**Figure S2: Crystal structure comparison of the eIF4A-(AG)5 complex clamping by a) RocA, b) silvestrol and c) DMPatA.** a-b) Phe163 is involved in π-π stacking interaction which rings B and C of RocA and silvestrol. Ring B is also engaged in additional π-π stacking interaction with RNA G8 and ring A with RNA A7. The two rocaglates are further stabilized in the RNA binding pocket by hydrogen-bonds with Gln195 and G8. b) Silvestrol has an additional dioxanyloxy-moiety which further interacts with the RNA residues A9 and G10 as well as with Arg110. c) DMPatA mimics the typical interactions of rocaglates except for the loss of interactions in pocket C. In addition, it is involved in a hydrogen bond with Asp198.

**Figure S3: Cloning scheme of insect-adapted dual-luciferase vectors** to analyse the effect of silvestrol on translation efficiency. The first step was the replacement of the HSV-TK promoter with the hr5-ie1 enhancer-promoter. In the second step, the Type III Hepatitis C virus (HCV) internal ribosome entry site was replaced with a Type IV *Drosophila* C virus IRES. Translation of both IRES types is eIF4A1-independent (33).

## References

1. Agboli E, Zahouli JB, Badolo A, Jöst H. Mosquito-associated viruses and their related mosquitoes in West Africa. Viruses. 2021;13(5):891.

2. Chu J, Galicia-Vázquez G, Cencic R, Mills JR, Katigbak A, Porco JA, et al. CRISPR-mediated drug-target validation reveals selective pharmacological inhibition of the RNA helicase, eIF4A. Cell reports. 2016;15(11):2340–7.

3. Biedenkopf N, Lange-Grünweller K, Schulte FW, Weißer A, Müller C, Becker D, et al. The natural compound silvestrol is a potent inhibitor of Ebola virus replication. Antiviral research. 2017;137:76–81.

4. Elgner F, Sabino C, Basic M, Ploen D, Grünweller A, Hildt E. Inhibition of Zika virus replication by silvestrol. Viruses. 2018;10(4):149.

5. Henss L, Scholz T, Grünweller A, Schnierle BS. Silvestrol inhibits chikungunya virus replication. Viruses. 2018;10(11):592.

6. Müller C, Obermann W, Karl N, Wendel H-G, Taroncher-Oldenburg G, Pleschka S, et al. The rocaglate CR-31-B (−) inhibits SARS-CoV-2 replication at non-cytotoxic, low nanomolar concentrations *in vitro* and *ex vivo*. Antiviral Research. 2021;186:105012.

7. Müller C, Obermann W, Schulte FW, Lange-Grünweller K, Oestereich L, Elgner F, et al. Comparison of broad-spectrum antiviral activities of the synthetic rocaglate CR-31-B (−) and the eIF4A-inhibitor Silvestrol. Antiviral Research. 2020;175:104706.

8. Müller C, Schulte FW, Lange-Grünweller K, Obermann W, Madhugiri R, Pleschka S, et al. Broad-spectrum antiviral activity of the eIF4A inhibitor silvestrol against corona-and picornaviruses. Antiviral research. 2018;150:123–9.

9. Taroncher-Oldenburg G, Müller C, Obermann W, Ziebuhr J, Hartmann RK, Grünweller A. Targeting the DEAD-box RNA helicase eIF4A with rocaglates—a pan-antiviral strategy for minimizing the impact of future RNA virus pandemics. Microorganisms. 2021;9(3):540.

10. Todt D, Moeller N, Praditya D, Kinast V, Friesland M, Engelmann M, et al. The natural compound silvestrol inhibits hepatitis E virus (HEV) replication *in vitro* and *in vivo*. Antiviral research. 2018;157:151–8.

11. Abdelkrim YZ, Harigua-Souiai E, Bassoumi-Jamoussi I, Barhoumi M, Banroques J, Essafi- Benkhadir K, et al. Enzymatic and molecular characterization of anti-*leishmania* molecules that differently target *leishmania* and mammalian eIF4A proteins, LieIF4A and eIF4A_Mus_. Molecules. 2022;27(18):5890.

12. Iyer KR, Whitesell L, Porco Jr JA, Henkel T, Brown LE, Robbins N, et al. Translation inhibition by rocaglates activates a species-specific cell death program in the emerging fungal pathogen *Candida auris*. MBio. 2020;11(2):10.1128/mbio. 03329–19.

13. Langlais D, Cencic R, Moradin N, Kennedy JM, Ayi K, Brown LE, et al. Rocaglates as dual-targeting agents for experimental cerebral malaria. Proceedings of the National Academy of Sciences. 2018;115(10):E2366–E75.

14. Obermann W, Azri MFD, Konopka L, Schmidt N, Magari F, Sherman J, et al. Broad anti-pathogen potential of DEAD box RNA helicase eIF4A-targeting rocaglates. Scientific Reports. 2023;13(1):9297.

15. Bryden SR, Pingen M, Lefteri DA, Miltenburg J, Delang L, Jacobs S, et al. Pan-viral protection against arboviruses by activating skin macrophages at the inoculation site. Science Translational Medicine. 2020;12(527):eaax2421.

16. Rosales-Rosas AL, Goossens S, Chiu W, Majumder A, Soto A, Masyn S, et al. The antiviral JNJ- A07 significantly reduces dengue virus transmission by *Aedes aegypti* mosquitoes when delivered via blood-feeding. Science Advances. 2024;10(48):eadr8338.

17. Iwasaki S, Floor SN, Ingolia NT. Rocaglates convert DEAD-box protein eIF4A into a sequence-selective translational repressor. Nature. 2016;534(7608):558–61.

18. Pestova TV, Kolupaeva VG, Lomakin IB, Pilipenko EV, Shatsky IN, Agol VI, et al. Molecular mechanisms of translation initiation in eukaryotes. Proceedings of the National Academy of Sciences. 2001;98(13):7029–36.

19. Iwasaki S, Iwasaki W, Takahashi M, Sakamoto A, Watanabe C, Shichino Y, et al. The translation inhibitor rocaglamide targets a bimolecular cavity between eIF4A and polypurine RNA. Molecular cell. 2019;73(4):738–48. e9.

20. Naineni SK, Bhatt G, Jiramongkolsiri E, Robert F, Cencic R, Huang S, et al. Protein–RNA interactions mediated by silvestrol—insight into a unique molecular clamp. Nucleic Acids Research. 2024;52(20):12701–11.

21. Magari F, Messner H, Salisch F, Schmelzle SM, van Zandbergen G, Fürstner A, et al. Potent anti-coronaviral activity of pateamines and new insights into their mode of action. Heliyon. 2024;10(13).

22. AAT Bioquest I. Quest graph™ IC50 calculator. ATT Bioquest, Inc.; 2024.

23. Wolfe AL, Singh K, Zhong Y, Drewe P, Rajasekhar VK, Sanghvi VR, et al. RNA G-quadruplexes cause eIF4A-dependent oncogene translation in cancer. Nature. 2014;513(7516):65–70.

24. Devignot S, Sha TW, Burkard TR, Schmerer P, Hagelkruys A, Mirazimi A, et al. Low-density lipoprotein receptor–related protein 1 (LRP1) as an auxiliary host factor for RNA viruses. Life Science Alliance. 2023;6(7).

25. Rodems SM, Friesen PD. Transcriptional enhancer activity of hr5 requires dual-palindrome half sites that mediate binding of a dimeric form of the baculovirus transregulator IE1. Journal of virology. 1995;69(9):5368–75.

26. Li J, Handler AM. Temperature-dependent sex-reversal by a *transformer-2* gene-edited mutation in the spotted wing drosophila, *Drosophila suzukii*. Scientific Reports. 2017;7(1):12363.

27. Cherry S, Doukas T, Armknecht S, Whelan S, Wang H, Sarnow P, et al. Genome-wide RNAi screen reveals a specific sensitivity of IRES-containing RNA viruses to host translation inhibition. Genes & development. 2005;19(4):445–52.

28. Lan Q, Fallon A. Small heat shock proteins distinguish between two mosquito species and confirm identity of their cell lines. American Journal of Tropical Medicine and Hygiene. 1990;43(6):669–76.

29. Schneider I. Cell lines derived from late embryonic stages of *Drosophila melanogaster*. Development. 1972;27(2):353–65.

30. Holm S. A simple sequentially rejective multiple test procedure. Scandinavian journal of statistics. 1979:65–70.

31. Šidák Z. Rectangular confidence regions for the means of multivariate normal distributions. Journal of the American statistical association. 1967;62(318):626–33.

32. Dunn OJ. Multiple comparisons using rank sums. Technometrics. 1964;6(3):241–52.

33. Svitkin YV, Siddiqui N, Sonenberg N. Protein synthesis initiation in eukaryotes: IRES-mediated internal initiation. eLS. 2015:1–11.

